# The microbiota of fresh unfermented date palm sap: a culturomics and metagenomics investigation

**DOI:** 10.1101/2025.01.10.632404

**Authors:** S. Djemal, M. Gdoura, N.E. Mathlouthi, R. Gdoura, C. Cassagne, H. Oumarou Hama

**Author notes:** These two authors equally participated in the work (S. Djemal, M. Gdoura). Corresponding author: Dr. Hamadou Oumarou Hama, MEPHI, IHU-Méditerranée Infection,19–21 Boulevard Jean Moulin, 13385 Marseille cedex 05, France,.

## Abstract

Palm sap is an artisanal fermented beverage, the consumption of which has been historically attested for millennia. Despite this, the underlying fermentative microbiota is poorly known. Here, three date palm sap samples collected and bought in Tunisia were investigated for bacteria, methanogenic archaea, and fungi using culturomics and molecular detection techniques, including metagenomics. Metagenomics yielded 250 bacteria, two fungi, and no archaea. Culturomics yielded 36 bacteria, eight fungi, and no archaea. Taken together, these approaches detected 250 bacteria and nine fungi, including 28 bacteria and eight fungi which had been previously unreported in date palm sap, in addition to three potentially new bacteria. Specifically, the bacteria *Zymomonas mobilis* and *Leuconostoc mesenteroides* and the fungus *Saccharomyces cerevisiae* were the most abundant, fermentative species found in the three samples, by culturomics and metagenomics. While determining the source of these microbes of interest was beyond the scope of the study, the data reported here may pave the way for a better understanding of the organic and organoleptic qualities of date palm sap in productive countries.

**IMPORTANCE:** This study provides valuable insights into the microbial composition of date palm sap, a traditional fermented beverage with deep historical roots. Despite its widespread consumption and cultural importance, the microbes driving its fermentation remain largely unexplored. Using advanced methods like metagenomics and culturomics, we uncovered a diverse array of bacteria and fungi, including species crucial for fermentation, such as *Zymomonas mobilis, Leuconostoc mesenteroides*, and *Saccharomyces cerevisiae*. Moreover, the discovery of previously unreported microbes highlights the potential for further research into their roles in sap fermentation and quality. Improving our understanding of the microbial communities involved in the fermentation of date palm sap underscores its importance to the field of applied and environmental microbiology.

## INTRODUCTION

Date palm sap is widely consumed under various local names in countries where the palm tree has been cultivated for millennia. While the first records of date palm sap mention it in Egypt, sap-derived wine was reported as the “drink of life” in an inscription dating from the reign of Merenre in the sixth dynasty, in Babylonia in 2600 BCE (1). This wine was shipped up the Nile as an Egyptian product destined for sub-Saharan regions of Africa.(1) Nowadays, this natural beverage is collected by tapping the tree to allow the liquid to flow into containers, often in the early morning to preserve its freshness and prevent excessive fermentation. Due to their abundance in sugars, amino acids, vitamins, and minerals, both xylem and phloem saps are harnessed for consumption as fresh or fermented drinks and as processed products including syrups, sugars, and sweeteners.(2) Notably, phloem saps from palms including the palmyra palm (*Borassus flabellifer*), the coconut palm (*Cocos nucifera* L.), the African oil palm (*Elaeis guineensis*), and the date palm (*Phoenix dactylifera* L.) are increasingly utilised to create a variety of value-added food products for human consumption.(3) In this situation, the alcohol in non-fresh palm sap most probably results from sugar fermentation, although the microorganisms responsible for such fermentation have not been reported.

The microbial composition of legmi (the local name for date palm sap in Tunisia where the present study was conducted), was previously examined through conventional culture-dependent techniques,(4) an approach which may capture less than 1% of microbial diversity due to inherent challenges in isolation methods.(5) The emergence of culture-independent molecular techniques has increased the ability to delve into microbial diversity at the metagenomic level, offering a more comprehensive understanding of the intricate microbial profile of legmi.

Our investigation aimed at uncovering the comprehensive microbial diversity in fresh, unfermented legmi, including less studied methanogenic archaea, with the potential to enhance our understanding of the microbial characteristics inherent in this beverage.

## MATERIALS AND METHODS

### Date palm sap samples

Three date palm sap samples were purchased in January 2024 from three different local producers located in the Gabes and Sfax regions, in Tunisia (Figure 1). In the Sfax region, one date palm sap sample was collected from a seven-metre-tall, female *P. dactylifera* L date palm tree (variety *Bou hettem*) located 15 km from Sfax city centre (34°49′08.8″N 10°38′59.8″E: E3 sample). In the Gabes region (Tbelbo), two date palm sap samples were collected from a three-metre-tall female *P. dactylifera* L tree, variety Agewi (33°50′09.2″N 10°06′57.3″E : E1 sample) and from a seven-metre-tall female *P. dactylifera* L tree, variety Temri (33°49′58.1″N 10°07′08.4″E: E2 sample). In each case, sap was collected using the non-destructive technique of tapping the palms. This was done early in the morning and during the initial days of the onset of production, avoiding direct exposure to sunlight. At the collection site, generally speaking the sap flows into non-sterile jugs set up by the farmer, who sells out the sap once he has retrieved the sap-filled jugs. Using a sterile syringe, one millilitre and three millilitres of sap were seeded, respectively and in ambient air in a sterile Hungate tube (Dominique Dutscher, Brumath, France) and a Falcon tube containing transport medium. Samples were placed in a well-sealed cooler containing dry ice until they were shipped at ambient temperature to the IHU-Méditerranée Infection in Marseille, France for further investigation, as presented below.

**Figure 1.**
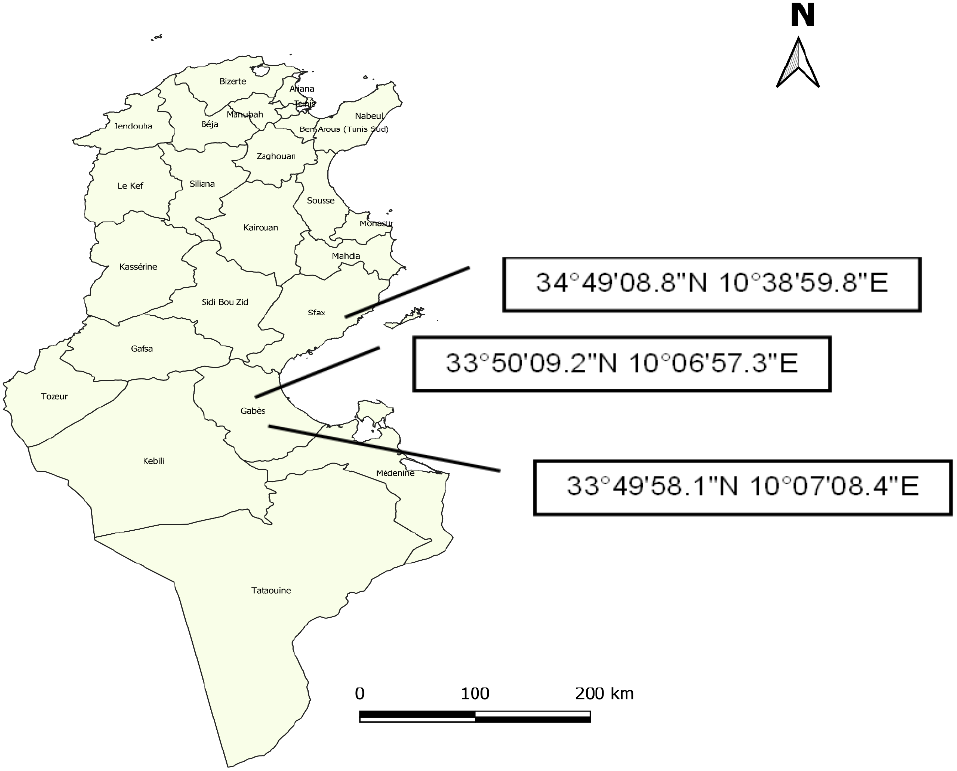
Location of samples E1: 33°50′09.2″N 10°06′57.3″E, E2: 33°49′58.1″N 10°07′08.4″E and E3: 34°49′08.8″N 10°38′59.8″E on the map of Tunisia.

### Legmi metagenomics

Total DNA extracted using the KingFisher automated extraction system and quantified using the Qubit™ dsDNA HS assay kit (Thermo Fisher Scientific, Eugene, Oregon USA), was sequenced on a MiSeq platform using a paired-end strategy after barcoding to be mixed with other genomic projects according to the Nextera XT library kit in a 2 × 250-bp format (Illumina Inc., San Diego, CA). The quality of the reads recovered was checked using FastQC software (version 0.11.9, https://www.bioinformatics.babraham.ac.uk/projects/fastqc) and data were cleaned using Trimmomatic software (Version 0.38). The taxonomic classification was performed using Kraken 2 software on the Bacterial and Viral Bioinformatics Resource Center online platform (BV-BRC, version 3.36.16: https://www.bv-brc.org/). The alpha and beta bacterial diversity indices in all the samples were compared using the Bray-Curtis index.

### Bacteria and methanogen culturomics

Bacterial culturomics was performed as previously described, with modifications.(6) One hundred microlitres of homogenised sap transport medium supernatant was inoculated into each of two BACT/ALERT bottles (bioMérieux, La Balme-les-Grottes, France): one designated for anaerobic culture and the other for aerobic culture, with follow-up at 1 day, 3 days, 7 days, 10 days, 15 days, 21 days, and 30 days. For each culture follow-up time, 1 mL of sample was collected from the BACT/ALERT bottle and serial dilutions were prepared from 1/10 to 1/10^10^ with Dulbecco’s phosphate buffer (DPBS). Each dilution was inoculated on 5% sheep blood agar plates (bioMérieux) incubated at 37 °C for 48 hours, one plate being incubated under atmospheric conditions and the other under an anaerobic atmosphere. As for methanogens, a 100 μL sample of sap was seeded in ambient air in a sterile Hungate tube (Dominique Dutscher, Brumath, France) containing 5 mL of GG medium, as previously described.(7, 8) Methanogen growth was monitored by detecting and quantifying methane by gas chromatography at D0 and D9, as previously described.(9)

All visible colonies were sub-cultured on the same culture medium they grew and were identified by matrix assisted laser desorption ionization-time of flight mass spectrometry (MALDI-TOF-MS) (Microflex, Bruker Daltonik, Bremen, Germany), as previously described.(10) Colonies with an identification score > 2 with a unique hit were considered as accurately identified at the species level. In other cases, (i.e., low score or multiple hits with same score), 16S rRNA gene sequencing was carried out to achieve the final identification.(11)

### Fungi culturomics

Date palm sap samples were diluted with DPBS and cultured on potato dextrose agar (PDA, Sigma-Aldrich, Saint-Quentin Fallavier, France), Sabouraud dextrose agar (BD diagnostic system, Le Pont-de-Claix, France), Dixon agar (Sigma-Aldrich) supplemented with 0.05 mg/mL chloramphenicol and 0.2 mg/mL cycloheximide and home-made FastFung medium, as previously described (12). Plates were incubated for 12 days at 28 °C in ambient atmosphere, and each yeast colony was sub-cultured on CHROMagar™ Candida media and identified by MALDI-TOF-MS, as previously described (13).

### Methanogen PCR

Total DNA was extracted from three samples (E1, E2, and E3) using an EZNA Tissue DNA kit (QIAGEN, Hilden, Germany). All samples were screened by standard polymerase chain-reaction (PCR) targeting the 16S rRNA archaeal gene with the primers SDArch0333aS15 (50-TCCAGGCCCTACGGG-30) and SDArch0958aA19 (50-YCCGGCGTTGAMTCCAATT-30) (Eurogentec, Seraing, Belgium), as previously described (Paul *et al*., 2004). In addition, real-time PCR assays specifically targeting the methanogen 16S rRNA gene were performed using the following primers and probes: Metha_16S_2_MBF (50-CGAACCGGATTAGATACCCG - 30); Metha_16S_2_MBR (50-CCCGCCAATTCCTT TAAGTT-30); and Metha_16S_2_MBP 6FAM-CCTGGGAAGTACGGTCGCAAG, as previously described (14). Master Mix and RNase-free water were used as negative controls.

## RESULTS

### Legmi microbiome: a metagenomics view

The mean quantity of DNA extracted from each sample was 17.9 ng/µL; 13.5 ng/µL and 1.41 ng/µL, for samples E1, E2, and E3, respectively, and no DNA was detected in the extraction blank used as a negative control. The sequencing of this DNA generated 1.1 Gb of data, with more than three million reads and an average of 1 331 030 reads per sample. After trimming and mapping against the BV-BRC database, 450 624; 257−541, and 916 312 bacterial reads were found, representing 35.7, 17.7, and 71.8% of all the reads generated in samples E1, E2, and E3, respectively. Taxonomic data generated by Kraken 2, normalised and filtered based on a low count and low variance filter, disclosed four bacterial phyla *Firmicutes, Actinobacteria, Proteobacteria*, and *Cyanobacteria* in all legmi samples, with *Firmicutes and Proteobacteria* being found in common, nine classes, 26 orders, 38 families, 78 genera, and 250 species in the three samples (Figure 2).

**Figure 2.**
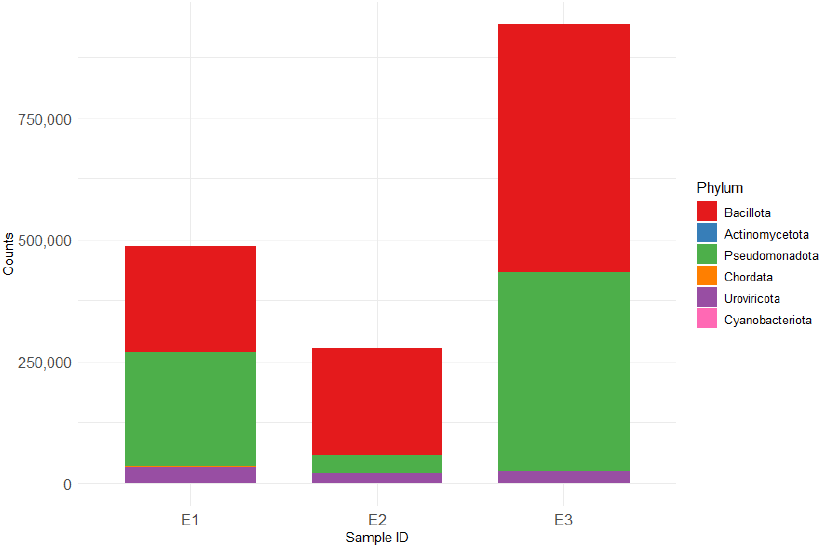
Bar plot representing relative abundance of detected phylum in fresh legmi.

The alpha and beta diversity indices (Figure 3) revealed that alpha diversity indicated a higher richness of bacterial genera in sample E1 compared to samples E2 and E3. In terms of beta diversity, the Bray-Curtis index revealed significant differences in the taxonomic compositions among the three sample sets. This measure, commonly used in ecological studies, quantifies the dissimilarity between two sets based on the abundance of species. In this case, the index highlighted that the species composition varied considerably across the E3 and E2 samples (0.81) and across the E1 and E2 samples (0.73), indicating that each sample contained distinct species. The Bray-Curtis index showed subtler differences in taxonomic composition between samples E3 and E1 (0.459). While some variation was still present, the dissimilarity between these two sets of samples was less pronounced compared to the others, suggesting a closer resemblance in their species composition.

**Figure 3.**
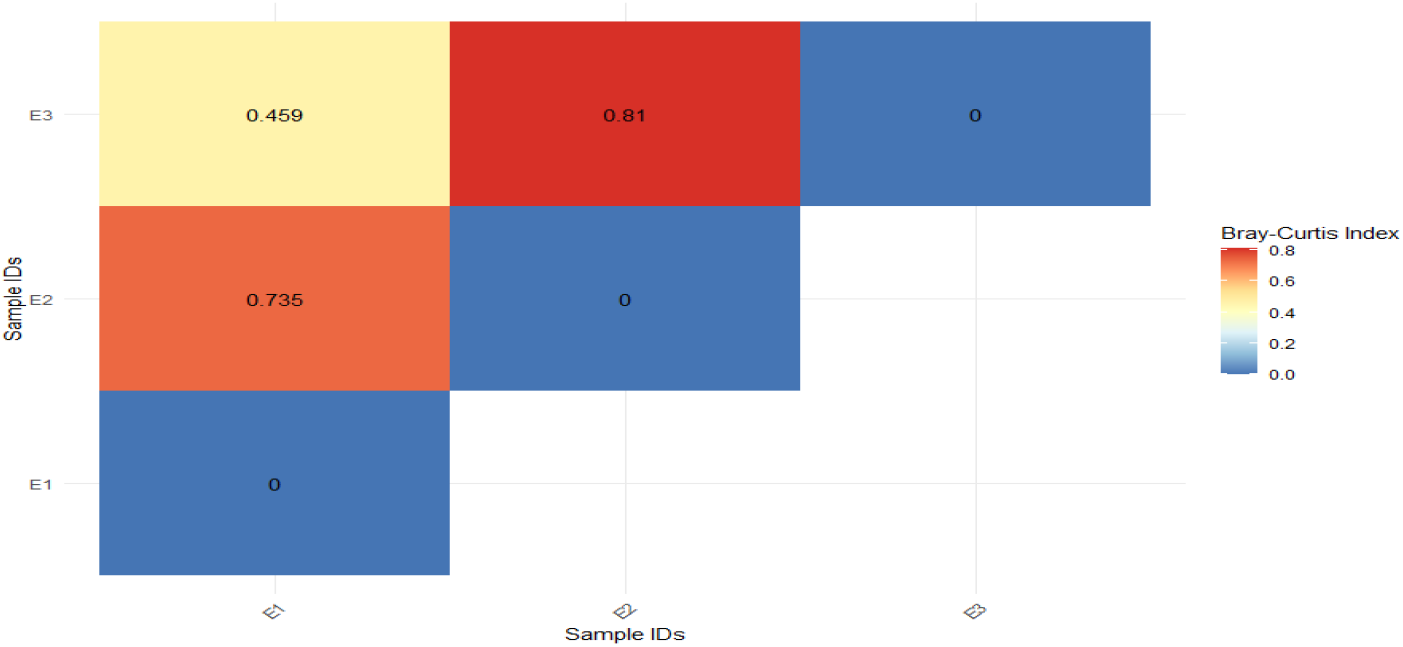
Bray-Curtis dissimilarity matrix for the bacterial communities in the three samples: E1, E2, and E3.

The bacterial composition of samples E1 and E2, both collected from the Gabes region, exhibited notable similarities and differences in terms of genus and species abundance. In both samples, the *Leuconostoc* genus dominates the microbial community: *Leuconostoc mesenteroides* (*L. mesenteroides*) was the most dominant species in E1, accounting for approximately 36.4% of total microbial composition (Supplementary Figure E1). Other notable species include *Rahnella aquatilis* (14.2%) and *Pseudomonas simiae* (7.7%).

*Leuconostoc citreum* and other *Leuconostoc* species, although present, show significantly lower abundance compared to *L. mesenteroides*. In E2, *L. mesenteroides* also dominates, representing a striking 52.8% of the bacterial population, further highlighting the shared microbial profile with E1 (Supplementary Figure E1). Other species such as *Leuconostoc suionicum* and *Leuconostoc pseudomesenteroides* were present in both samples, albeit in varying proportions, with *Leuconostoc* sp. also constituting a significant portion at 12.1% in E2. Similarly, the presence of other species such as *Pseudomonas putida* and *Rahenella aquatilis* was more pronounced in E2 (Supplementary Figure E2).

When comparing the bacterial profiles of E1 and E2 (both collected from the Gabes region) to E3 (collected from the Sfax region), *Zymomonas mobilis* (*Z. mobilis*) was the most abundant species (42.7%) in E3, representing a large proportion of the microbial community, reflecting the influence of geographic and environmental factors. Despite the dominance of *Leuconostoc* species in E1 and E2, in E3, *Leuconostoc sp*. still constituted a significant portion of the bacterial community, representing 24% of the population. However, the composition within the *Leuconostoc* genus varied. For instance, *Leuconostoc suionicum* was more abundant in E3 (14.7%) compared to its lower presence in E1 and E2. *L. mesenteroides*, although present, was less abundant in E3 (9.2%) than in E1 and E2, where it dominated. *Lactiplantibacillus plantarum* was highly abundant in E3 with over 40%, but much less common in E2 and E1, where its presence was minimal. The two species *Lactococcus lactis* and *Leuconostoc citreum* were more prominent in E1 than in the other samples, although still relatively low compared to other dominant species in E2 and E3 (Figure 4) (Supplementary Figure E3).

**Figure 4.**
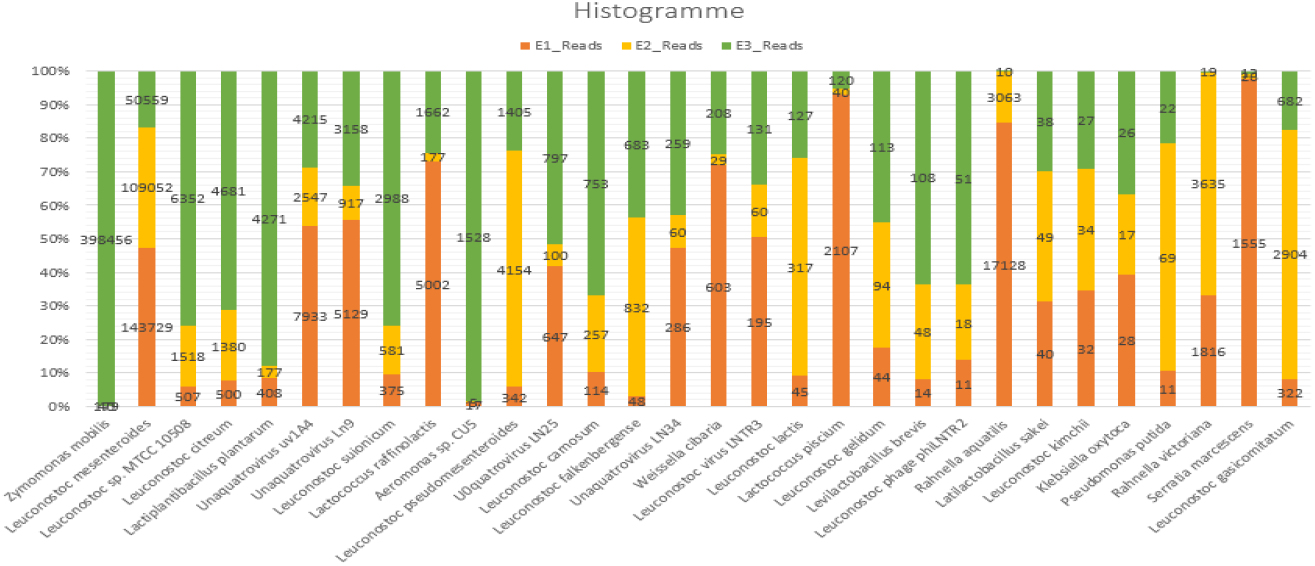
Histogram illustrates the relative abundance of different bacterial species across three sample sets: E1 (orange), E2 (yellow), and E3 (green). The Y-axis represents the percentage of abundance, while the X-axis lists the bacterial species identified.

### Legmi bacteriome: a culturomics view

Culturomics study of the three legmi samples yielded 36 bacteria identified by MALDI-TOF-MS, belonging to 17 genera, nine families, and three phyla: *Actinobacteria, Firmicutes*, and *Proteobacteria*. Among these phyla, *Firmicutes* were the most prevalent. Analysis of the bacterial species identified in three samples (E1, E2, and E3) using a Venn diagram showed that sample E1 contained six exclusive species, while samples E2 and E3 contained eight and 10 exclusive species, respectively. The intersection between E1 and E2 contained two species, and the intersection between E2 and E3 contained three species. Notably, the intersection between all three samples E1, E2, and E3 included seven shared species. Interestingly, no species were shared exclusively between E1 and E3, as indicated in Figure 5. In addition, four isolates remained unidentified after careful MALDI-TOF-MS analysis: one cultured from E1, two from E2, and one from E3. These four isolates, deposited in the Souches de l’Unité des Rickettsies (CSUR) collection under CSURQA0360, CSURQA0359, CSURQA0358 and CSURQA0357, will undergo further complementary investigations to confirm the degree of novelty of their identification.

**Figure 5.**
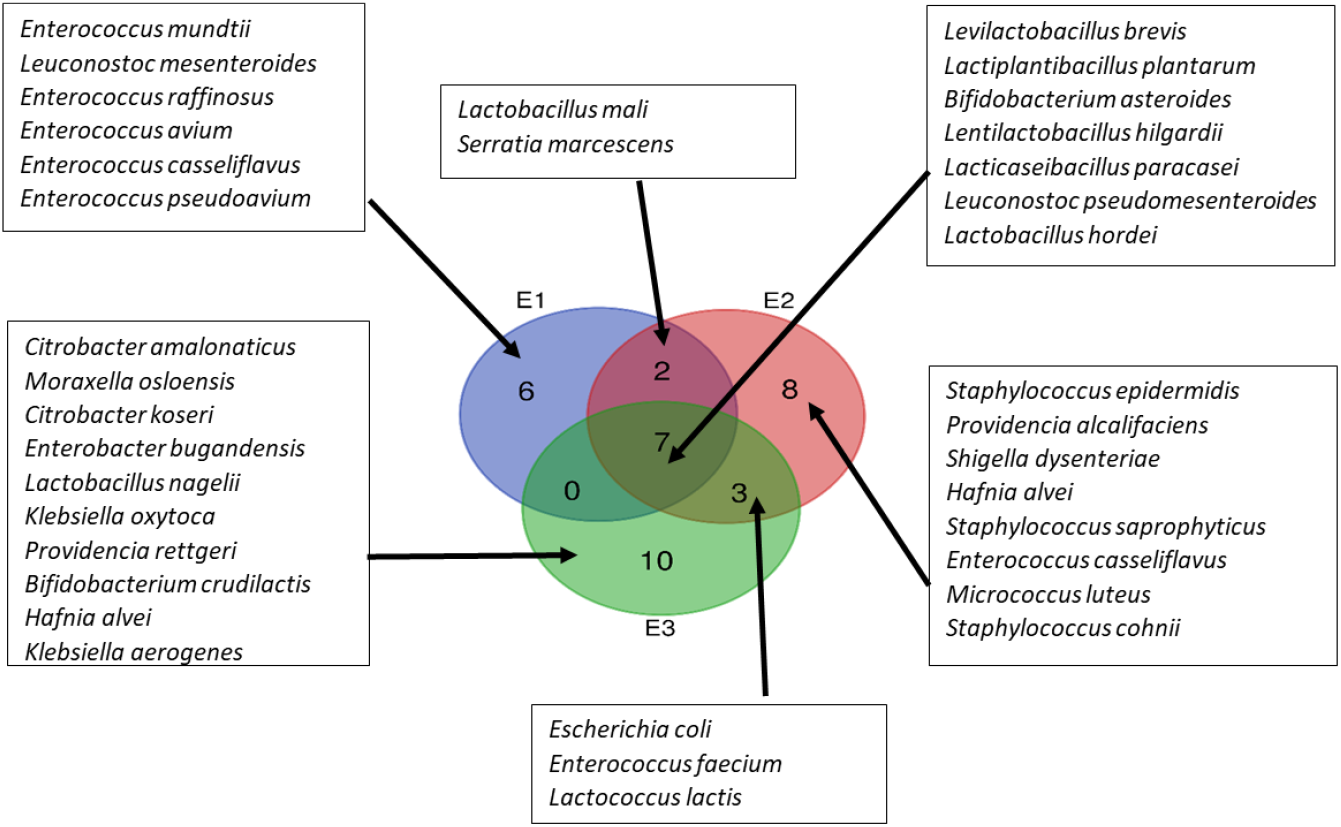
Venn diagram representing species isolated based on bacterial culturomics from three legmi samples.

### Legmi fungome: a culturomics view

Two common fungal species, *Saccharomyces cerevisiae* and *Hanseniaspora valbyensis*, were cultured in the three samples E1, E2, and E3. In addition, *Candida krusei* and *Pichia kudriavzevii* were identified in samples E2 and E3, indicating some continuity in yeast diversity between these two samples, which was absent in sample E1. Specifically, sample E1 was distinguished by the presence of *Hanseniaspora guilliermondii* and *Lachancea thermotolerans*, while sample E3 was distinguished by *Kazachstania exigua* and *Kazachstania humilis*. These findings highlight significant variations in yeast composition between the samples (Figure 6).

**Figure 6.**
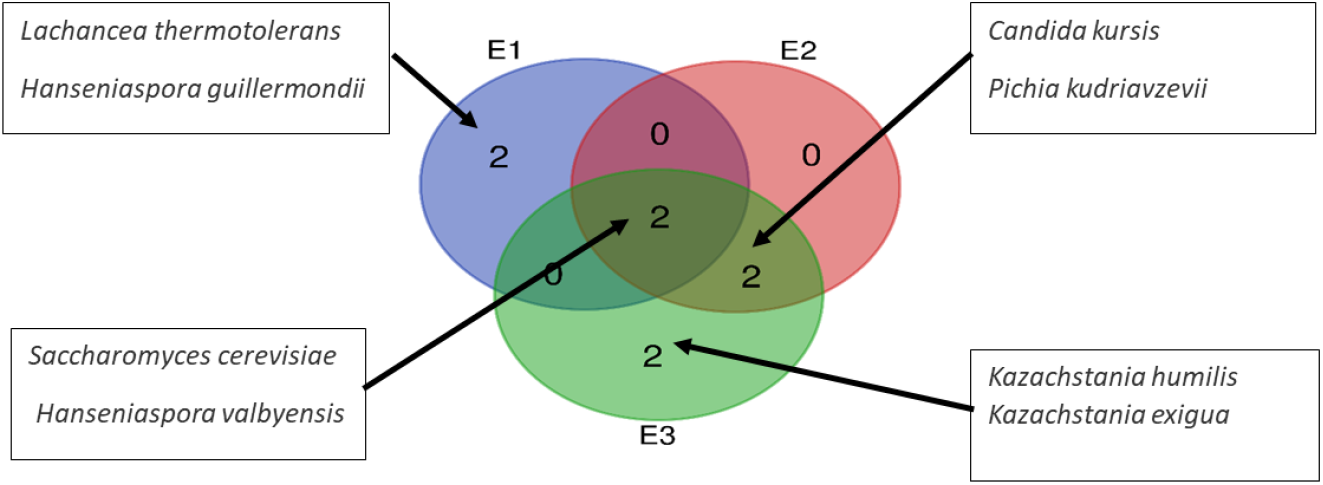
Venn diagram representing species isolated based on fungi culturomics from three legmi samples.

### Legmi methanodrome: a combined view

In this study, the three legmi samples (E1, E2, and E3) were analysed for the presence of methanogenic archaea using real-time PCR and PCR sequencing targeting the broad-range archaeal 16S rRNA gene. While real-time PCR yielded positive results for the presence of methanogens in sample E2, neither E1 nor E3 exhibited detectable methanogens through this technique with negative controls remaining negative. Furthermore, PCR sequencing did not reveal any methanogens across all samples, and the detection of methane in these cultured samples was negative.

## DISCUSSION

Using complementary culturomics and metagenomics methods, we analysed the microbial diversity of three legmi samples. The legmi microbiota were investigated in the presence of negative controls to validate the data. It should, however, be borne in mind that this may not reflect the actual composition of the palm juice itself, as it had been collected from producers using non-sterile cans. Combining culturomics with metagenomics allowed for a thorough exploration of microbial diversity. Culturomics facilitated the isolation and characterisation of specific bacterial strains, while metagenomics provided insights into the broader microbial community including microbial networks.

We confirmed previously reported bacterial and fungal repertoires of legmi. Of 19 genera of bacteria and 14 genera of fungi previously reported in four palm sap types (oil palm: *Elaeis guineensis*, raffia palm: *Raphia hookeri*, date palm: *P. dactylifera* L, coconut palm: *Cocos nucifera*, and ron palm: *Borassus aethiopum*),(5, 15–19) 15 bacteria genera and six fungal genera were also detected in the present study, while 63 bacteria and one fungus were newly detected. In particular, *L. mesenteroides*, which was found to dominate across all three samples here investigated, is known to kick-off fermentation processes.(20) Bacteria such as *Leuconostoc* spp., *Lactobacillus* spp., and *Enterococcus* spp. also found in legmi, may offer health benefits, as these bacteria exhibit excellent probiotic characteristics, including antibiotic properties and hydroxyl radical-scavenging abilities (21).

Interestingly, we observed differences in microbial diversity among the three samples here investigated. *Rahnella aquatilis* was abundant in E1 (14.2%) and E2 (6.3%) but was absent in E3. *Rahnella* is typically associated with plant environments, which could indicate local contamination or plant-microbe interactions (22). In contrast, *Zymomonas mobilis* (*Z. mobilis*) was abundant in E3 but not in E1 or E2. *Z. mobilis* is a well-known fermentative bacterium, often associated with sugar-rich environments such as date sap (23). Such variations in bacterial species across the samples could be attributed not only to regional environmental conditions (E1 and E2 were collected from the same region of Gabes while E3 was collected in the region of Sfax) as factors such as climate, soil, and agricultural practices may contribute to variations in microbial populations, but the early fermentation processes initiated during time lapse between collection and analysis may also have influenced the detected repertoire of bacteria and fungi, due to the alcohol-susceptibility of most microbes.

Fermentation could have been initiated by dominant fungal species, as *Saccharomyces cerevisiae, Pichia kudriavzevii*, and *Lachancea thermotolerans*. All are known to thrive in high-sugar environments and to be commonly involved in the natural fermentation of fruit and sap (24). In addition, it should be noted that *S. cerevisiae* and *Hanseniaspora valbyensis* were the sole two species recovered from the three samples. This two-yeast combination is already used in several beverages based on fermentation, including kombucha (black tea-based) (25– 27) and cider (apple-based) to increase the production of esters, which are the basis of the pleasant, floral and fruity aroma of these drinks (28). Moreover, the presence in the E3 sample of additional yeast species, such as *Hanseniaspora guillermondii, Kazachstania exigua, Kazachstania taniahumilis*, and *Candida kursis*, might also be explained by different fermentation dynamics occurring during the time lag. Utilising Illumina MiSeq Pair End DNA sequencing, the analysis aimed to provide a comprehensive understanding of the microbial community present in this traditional beverage.

This research marks a significant contribution to the field, potentially representing the first comprehensive study focused on the microbiota of palm date sap. The findings could pave the way for future investigations into not only the fermentation processes for optimising fermentation conditions and improving product quality but also any health benefits associated with the consumption of this artisanal beverage.

## LINKS OF INTEREST

None to be declared.

## ACKNOWLEDKEMTS

The authors acknowledge the contribution of Prof. Michel Drancourt, Aix-Marseille Université, Marseille, France for study design and drafting.

